# Data-driven spatial filtering for improved measurement of cortical tracking of multiple representations of speech

**DOI:** 10.1101/551218

**Authors:** D Lesenfants, J Vanthornhout, E Verschueren, T Francart

## Abstract

**Objective:** Measurement of the cortical tracking of continuous speech from electroencephalography (EEG) recordings using a forward model is an important tool in auditory neuroscience. Usually the stimulus is represented by its temporal envelope. Recently, the phonetic representation of speech was successfully introduced in English. We aim to show that the EEG prediction from phoneme-related speech features is possible in Dutch. The method requires a manual channel selection based on visual inspection or prior knowledge to obtain a summary measure of cortical tracking. We evaluate a method to (1) remove non-stimulus-related activity from the EEG signals to be predicted, and (2) automatically select the channels of interest.

**Approach:** Eighteen participants listened to a Flemish story, while their EEG was recorded. Subject-specific and grand-average temporal response functions were determined between the EEG activity in different frequency bands and several stimulus features: the envelope, spectrogram, phonemes, phonetic features or a combination. The temporal response functions were used to predict EEG from the stimulus, and the predicted was compared with the recorded EEG, yielding a measure of cortical tracking of stimulus features. A spatial filter was calculated based on the generalized eigenvalue decomposition (GEVD), and the effect on EEG prediction accuracy was determined.

**Main results:** A model including both low- and high-level speech representations was able to better predict the brain responses to the speech than a model only including low-level features. The inclusion of a GEVD-based spatial filter in the model increased the prediction accuracy of cortical responses to each speech feature at both single-subject (270% improvement) and group-level (310%).

**Significance:** We showed that the inclusion of acoustical and phonetic speech information and the addition of a data-driven spatial filter allow improved modelling of the relationship between the speech and its brain responses and offer an automatic channel selection.

## 1. Introduction

Understanding the brain processing of speech could impact our knowledge of hearing disorders, such as sensorineural hearing loss, and allows the improvement of current hearing aids and cochlear implants. In particular, brain-based decoding of speech features could open doors to objective evaluation of a patient’s speech understanding. However, while time-locked event-related brain responses to an isolated acoustic event can be extracted by averaging over several presentations, the response of the brain to continuous natural speech is complex because of the overlapping responses of consecutive auditory tokens, the integration of multiple prosodic features dynamically changing over the continuous speech and, higher-order processes such as semantic analysis.

Low-frequency temporal fluctuations of the speech stimulus, i.e. the amplitude of the speech envelope, contain crucial information such as the speech rate (Ríos-López, Molnar, Lizarazu, & Lallier, 2017), and contribute to the perception of speech (Abrams, Nicol, Zecker, & Kraus, 2008; Giraud & Poeppel, 2012; Rosen, 1992; Shannon, Zeng, Kamath, Wygonski, & Ekelid, 1995). Therefore, the envelope has been widely used as an input of a linear model (see (Crosse, Di Liberto, Bednar, & Lalor, 2016) for more details) to predict the electroencephalographic (EEG) responses to speech. This linear model is called a forward model (Di Liberto & Lalor, 2016; Di Liberto, O’Sullivan, & Lalor, 2015). The inverse, a model (or decoder) to reconstruct the speech from the associated EEG recording, is called a backward model (N. Ding & Simon, 2012, 2013; O’Sullivan et al., 2015; Vanthornhout, Decruy, Wouters, Simon, & Francart, 2018). Increasingly complex attributes of speech are processed hierarchically in the brain, from simple acoustic deviants in the earlier areas, to low-level acoustic information and discrete features of speech (e.g., phonemes, syllables, words) in later areas (Chang et al., 2010; Okada et al., 2010; Peelle, Johnsrude, & Davis, 2010). Interestingly, Di Liberto et al. (2015, 2017) showed that the relationship between continuous English speech and its brain response could be best described using a model based on both the low-level acoustic (i.e., the spectrogram of the speech) and high-level discrete (i.e., phonetic tokens) representations (Di Liberto & Lalor, 2017; Di Liberto et al., 2015; Lesenfants, Vanthornhout, Verschueren, Decruy, & Francart, 2019).

Both the backward and the forward model have their limitations. A backward model is a data-driven approach for optimally combining time-shifted EEG channels: a range of delays (classically between 0 and 250 ms) is first applied to each channel, then all the delayed channels are weighted in order to linearly reconstruct the speech envelope. Inversely, a forward model combines the time-shifted speech envelope in order to predict the EEG responses to the speech stimulus. While the forward model can predict responses for individual EEG channels, allowing to study the topology of speech activation across the scalp, it requires a manual channel selection based on visual inspection or prior knowledge to obtain a summary measure of cortical tracking. On the other hand, the backward model will automatically weigh each channel’s relevance using the input data with the loss of spatial information, but with higher resulting reconstruction accuracies than the forward model. Moreover, the backward model requires continuous speech features, excluding its applicability to discrete speech features such as words or phonemes.

In the current study, we apply a data-driven method to remove non-stimulus-related activity from the EEG signals and automatically select the channels of interest, which is described in detail by Das, Vanthornhout, Francart, & Bertrand (2019). This method makes use of the Generalized Eigenvalue Decomposition GEVD (Hassani, Bertrand, & Moonen, 2016), which has been used to remove artefacts from the EEG (Somers, Francart, & Bertrand, 2018), and cancel EEG noise (Hajipour Sardouie, Shamsollahi, Albera, & Merlet, 2015; Serizel, Moonen, Van Dijk, & Wouters, 2014). It is closely related to the CCA-based method proposed by Dmochowski, Ki, DeGuzman, Sajda, & Parra (2018). Decomposing the stimulus-response relationship into multiple independent dimensions and spatially filtering by weighting channels’ relevance has allowed to efficiently capture the neural representations of speech stimuli in audio and audio-visual paradigms (de Cheveigné & Parra, 2014; Dmochowski, Ki, DeGuzman, Sajda, & Parra, 2018).

We here aim to (1) show that the temporal response function estimation and EEG prediction based on phonemes can be replicated in Dutch, (2) improve EEG prediction to several speech representations by removing non-stimulus-related components from the EEG signals, and (3) show that the channel selection problem of the forward model can be solved in a data-driven way using GEVD.

## 2. Methods

### 2.1 Participants

Eighteen Flemish-speaking volunteers (7 men; age 24 ± 2 years; two were left-handed) participated in this study. Each participant had normal hearing, verified by pure tone audiometry (thresholds lower than 20 dB HL for 125 Hz to 8000 Hz using a MADSEN Orbiter 922–2 audiometer). The study was approved by the Medical Ethics Committee UZ KU Leuven/Research (KU Leuven, Belgium) with reference S59040 and all participants provided informed consent.

### 2.2 Experiment

Each participant sat in an electromagnetically shielded and soundproofed room and listened to the children’s story “Milan”, written and narrated in Flemish by Stijn Vranken. “Milan” is a Dutch children’s story, similar to “Alice in Wonderland” for the Anglo-Saxon community. However, the Milan story is generally unknown. Therefore, we hypothesize that the participant had maximal attention during its listening and that the familiarity of the participant with this story should not affect the cortical tracking to the speech features. The stimulus was 14 minutes long and was presented binaurally at 60 dBA. It was presented through Etymotic ER-3A insert phones (Etymotic Research, Inc., IL, USA) which were electromagnetically shielded using CFL2 boxes from Perancea Ltd. (London, UK). The acoustic system was calibrated using a 2-cm^3^ coupler of the artificial ear (Brüel & Kjær, type 4192, Nærum, Denmark). The experimenter sat outside the room and presented the stimuli using the APEX 3 (version 3.1) software platform developed at ExpORL (Dept. Neurosciences, KU Leuven, Belgium) [(Francart, van Wieringen, & Wouters, 2008) and an RME Multiface II sound card (RME, Haimhausen, Germany) connected to a laptop running Windows.

### 2.3 Recordings

EEG signals were recorded from 64 Ag/AgCl ring electrodes at a sampling frequency of 8192 Hz using a Biosemi ActiveTwo system (Amsterdam, Netherlands). The electrodes were placed over the scalp according to international 10-20 standards.

### 2.4 Data analysis

All analyses were done with custom-made Matlab scripts and the mTRF Toolbox (Crosse et al., 2016; Gonçalves, Whelan, Foxe, & Lalor, 2014).

#### Speech features

We extracted five different representations of the speech stimulus, selected according to (Di Liberto et al., 2015):

1. The broadband amplitude envelope (Env) was extracted as the absolute value of the (complex) Hilbert transform of the speech signal.
2. The spectrogram representation (Sgram) was obtained by first filtering the speech stimulus into 16 logarithmically-spaced speech frequency bands using zero phase Butterworth filters with 80 dB attenuation at 10 % outside the passband between 250 Hz and 8 kHz, according to Greenwood’s equation (Greenwood, 1961), assuming linear spacing in the cochlea. We then calculated the energy in each frequency band using a Hilbert transform (see Env).
3. The time-aligned sequence of phonemes (Ph) was extracted using the speech alignment component of the reading tutor (Duchateau et al., 2009), which allows for reading miscues (skipped, repeated, misread words), automatically segmenting each word into phonemes from the Dutch International Phonetic Alphabet (IPA) and performing forced alignment. We then converted this into a binary matrix mask representing the starting and ending time-points (i.e., a ‘1’ from the start until the end of each presentation) for each 37 phonemes present in both the story and the matrix sentences.
4. The time-aligned binary sequence of phonetic features (Fea) was assembled using the following groups of phonemes: short vowels, long vowels, fricative consonants, nasal consonants and plosive consonants.
5. The combination of time-aligned sequence of phonetic features and the spectrogram as proposed by Di Liberto et al. (2015) (Di Liberto et al., 2015), hereinafter named FS.

#### EEG signal processing

EEG artifacts were removed using a multi-channel Wiener filter algorithm (Somers et al., 2018). We then re-referenced each EEG signal to a common-average reference before downsampling from 8192 to 1024 Hz to decrease processing time. EEG signals were bandpass filtered between 0.5-4 Hz (delta), 4-8 Hz (theta), 8-15 Hz (alpha), 15-30 Hz (beta), or 30-45 Hz (low gamma; for this frequency, we then computed the envelope of the filtered signals) using zero phase Butterworth filters with 80 dB attenuation at 10 % outside the passband. Stimulus representations and EEG signals were then downsampled to 128 Hz.

#### Temporal response function (TRF) estimation

Stimulus representation and EEG signals were then downsampled to 128 Hz. Each 14-min dataset was split into 7 consecutive non-overlapping 2-min segments (segment 1: from second 0 to second 120; segment 2: from second 120 to second 240; etc). A 7-fold leave-one-out cross-validation was used to separate the segments in a training set and a testing set. A quantitative mapping (i.e., a model) between each speech representation and the recorded EEG (training set) was computed based on a least-squares linear regression with ridge regression and a temporal integration window of 0-400 ms. As suggested by Crosse et al. (2016), we selected the value of the ridge regression regularization parameter by cross-validation. This mapping was used to predict EEG signals using each stimulus representation from the test set. The cortical speech tracking was computed as the correlation (Spearman’s rank-order correlation) between the recorded and predicted EEG signals at the different electrode locations. We calculated the averaged correlation over an area by first computing each electrode’s correlation and then averaging these correlations over the area.

A model was trained using either single-subject specific training data (ss or ssGEVDss; see panel A and C, Fig. 1) or a group-level training dataset (gen, genGEVDss or genGEVDgen; see panel B, D and E, Fig. 1). In the second case, the grand-average model is computed by averaging the covariance matrix across subjects. Averaging data across subjects (“grand-average”) has the benefit that more data is available, and subject-specific quirks are avoided, resulting in a more stable model. However, it has the downside that individual differences in brain anatomy and physiology are not taken into account. The different models specified below, are designed to leverage the strengths of both grand average and subject-specific models.

**Fig. 1.**
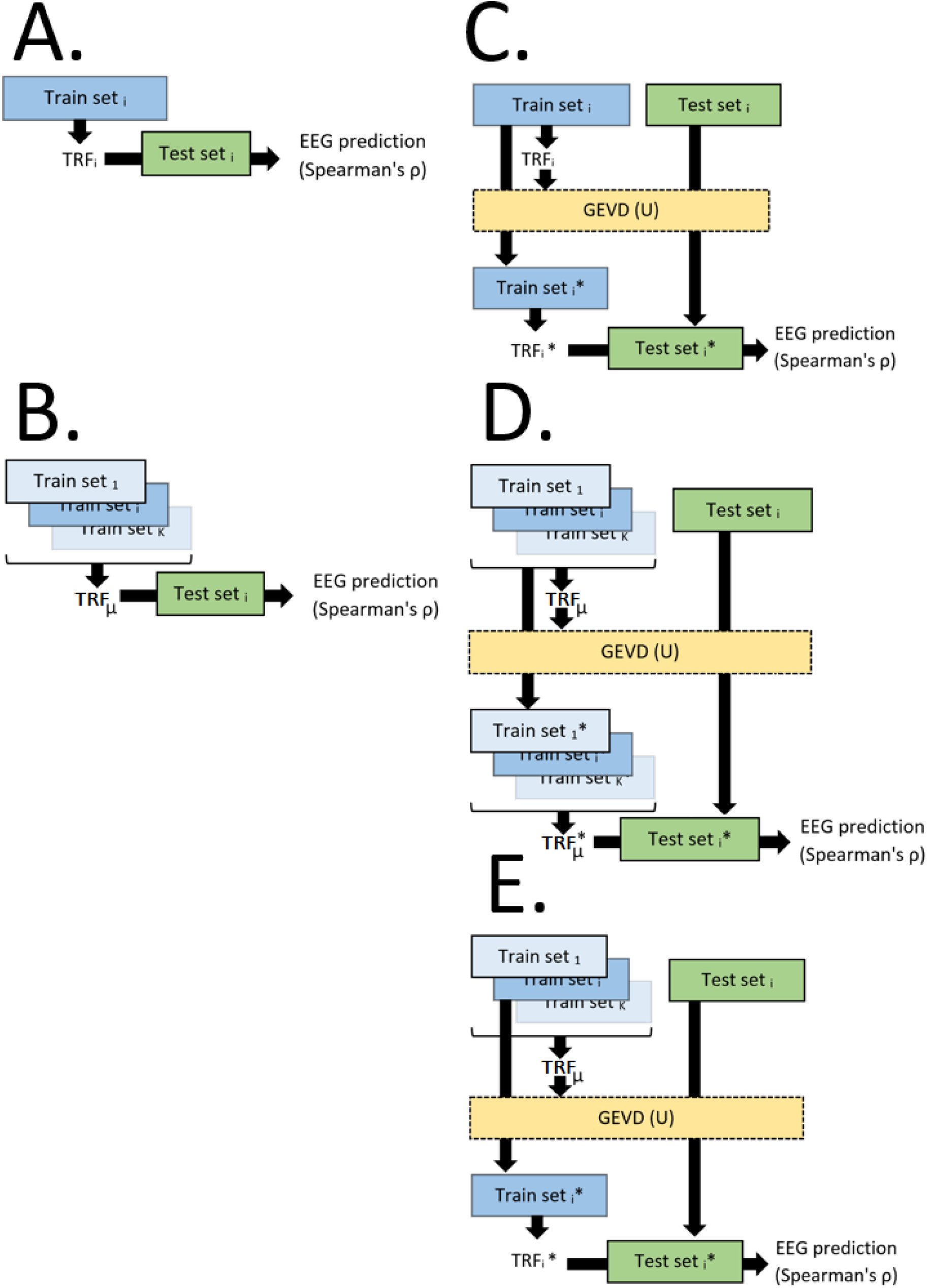
Illustration of the different algorithms evaluated in the current manuscript. A quantitative mapping (TRF) computed between the training speech and recorded EEG was used to predict EEG signals using each stimulus representation from the test set (i. The cortical speech tracking was computed as the Spearman’s correlation between the recorded and predicted EEGsignals at the different electrode locations. A model was trained using either single-subject specific training dataset (TRF_i_; see ss and ssGEVDss models in panels A and C respectively) or group-level training dataset (μTRF; see gen, genGEVDss and genGEVDgen models in panels B, D and E respectively). In ssGEVDss, genGEVDss and genGEVDgen models (see panels C, D and E), a spatial filter U was extracted from the training dataset to remove non-stimulus-related activity from the EEG signals by optimally mapping the recorded and predicted EEG signals. This spatial filter was finally applied to both training and testing dataset prior to rebuilding the TRF and computing the model’s performance. A set (or dataset) refers to the EEG data plus the corresponding stimulus features.

#### Spatial filtering

In the xGEVDy models (x = dataset used for training the initial TRF, y = GEVD-corrected dataset used for updating the TRF; x and y can either be a subject-specific, ss, or grand-averaged, gen, dataset), we applied a spatial filter **U** on the training and testing data allowing to remove non-stimulus-related activity from the EEG signals to be predicted by optimizing the mapping between the recorded and the predicted EEG signals from the training dataset. In other word, we here make the assumption that an EEG recording *e*(*t*) = *e_s_*(*t*) + *e_r_*(*t*) can be decomposed in EEG activity in response to the speech **e**_s_(*t*) and EEG activity unrelated to the speech **e**_r_(t), such that *U*[*e*(*t*)] = *U*[*e_s_*(*t*) + *e_r_*(*t*)] = *e_s_*(*t*). The optimization process in this method relies on the GEVD, which find linear combinations that maximise the SNR (based on predicted clean EEG and actual EEG). Considering an N-channel EEG recording *e*(*t*) ∈ *R^N^* and a N-channel EEG prediction *p*(*t*) ∈ *R^N^* based on a speech feature. The underlying assumption of this method is that a group of eigenvectors V can be extracted such that *R_E_* * *V* = *R_P_ * V * D*, with **R_E_**, the covariance matrix of the EEG recording = *E*{*ee^T^*}; **R_P_**, the covariance matrix of the EEG prediction *E*{*pp^T^*} and **D** the eigen-values. Computing this joint diagonalization of the covariance matrices allows to extract M components (with M ≤ N) associated with the M highest eigen-values such that the N-M speech-unrelated components can be filtered out the EEG recordings. The mapping **U** is then equal to the M first components of **D**, the remaining ones being set to zero.

A scheme of each model estimation is shown in Fig. 1. In ss model, TRFs were estimated using subject specific training data, then used to predict the EEG responses to the test speech data (see Fig. 1, panel A). In the gen model, the training datasets of all the participants were concatenated and used to extract the grand-average TRF of this group. Each averaged model is then convolved with data from the corresponding individual test set to predict the cortical responses to speech (see Fig. 1, panel B). In the ssGEVDss model, we first computed the subject-specific TRF as explained in the ss model. We then computed the mapping between the recorded training EEG and the predicted training EEG based on the speech stimuli of the training dataset. We then recomputed new TRFs based the newly mapped EEG training data (i.e., by applying the M-filtered-out-component eigenvectors of the GEVD on the recorded EEG training dataset related to the M-highest eigenvalues) and applied this corrected TRF to the mapped test set (see Fig. 1, panel C). The genGEVDgen relied on the same principle of the ssGEVDss except that we used the concatenated group training dataset rather than the subject-specific dataset in order to extract both the TRF and the spatial GEVD-mapping (see Fig. 1, panel D). Note that in the GEVD-based models, we can either back-project the M-selected components in the electrode space or compute correlations between components extracted from the recorded and predicted test set. In the following, to facilitate the visualization of areas related to speech activation, we will back-project the selected components in the electrode space.

#### Statistical analysis

A permutation test (Nichols & Holmes, 2002; Noirhomme et al., 2014) was used to evaluate the significance level for each model (1000 repetitions, p < .01). The significance of change between conditions was assessed with a non-parametric Wilcoxon signed-rank test (alpha = 0.01), with Bonferroni correction for multiple comparisons.

## 3. Results

In Fig 2, grand-average TRFs are shown for the different stimulus features, for a selection of fronto-temporal electrodes (F, Fc, Ft, C and T electrodes), using signal processing scheme A from Fig 1. The grand-average TRF of the model based on the envelope showed a positive activation at around 50 ms, 150 ms and 250 ms, and a negative activation at around 100ms (see Fig. 2, upper left panel). For the model based on the spectrogram (see Fig. 2, lower left panel), we observed a positive activation at early latencies and between 100 ms and 200 ms, as well as a negative activation at around 100 ms for frequencies below 600Hz. Frequencies above 600 Hz showed a positive activation at around 100 ms and a negative activation at around 50 ms. The phoneme-based model showed a positive activation for all the phonemes at 175ms (see Fig. 2, upper right panel) and a negative activation at around 125 ms. A majority of the phonemes (i.e., mainly the consonants) also showed an earlier positive activation at around 75 ms. All the different phonetic features showed a positive activation at 175 ms. Interestingly, while plosive (resp. the nasal and fricative) consonants showed a strong (resp. weak) activation at around 75 ms, the long vowels showed no earlier activation. Short vowels presented an activation at around 50 ms (see Fig. 2, lower right panel).

**Fig. 2.**
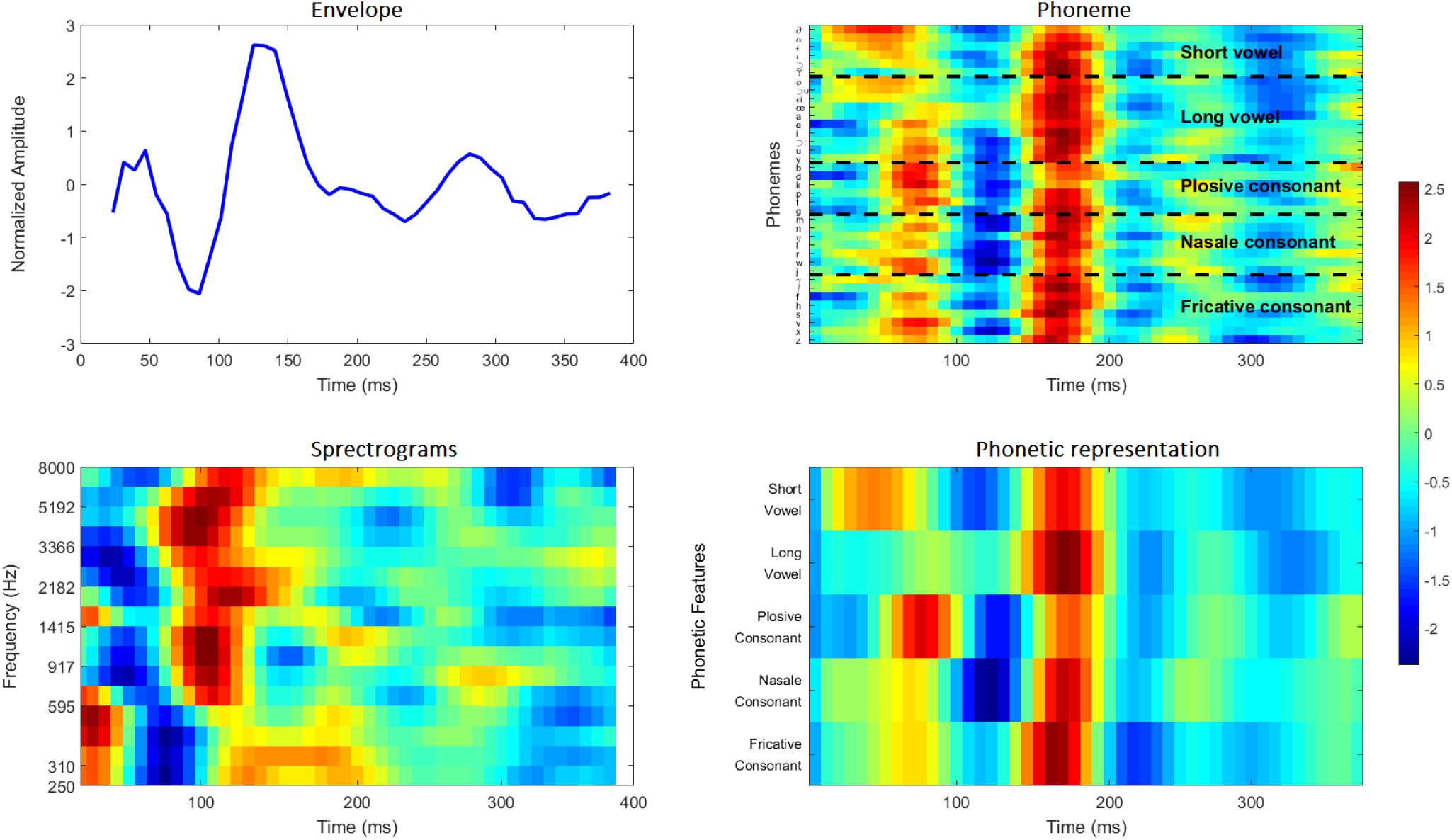
Grand-average Temporal Response Function for a model based on the envelope, the spectrogram, phoneme and phonetic representation averaged over cross-validation folds and fronto-temporal electrodes.

Fig. 3 is a replication of Di Liberto et al (2015) showing the impact of the frequency bands on the tracking of the speech features. Note that we decided to include more electrodes (see Fig. 3, lower right panel) in order to take into account topographical differences observed between the different frequency bands (see below; Fig. 4). The averaged EEG prediction in the frontotemporal regions decreased with the EEG frequency band frequency using a generic model. Using the delta or theta EEG band, the models based on FS showed higher EEG predictions than the others (Wilcoxon Signed-Rank Test, WSRT, p = 0.025 for Ph; p < 0.01 for Sgram and Env), except the Fea model (WSRT; for the delta band, p = 0.14, for the theta band, p = 0.73). The Fea-based model showed higher EEG predictions than the models based on either Env or Sgram (WSRT, using the delta band: p = 0.02 for Sgram, p < 0.01 for Env; using the theta band, p < 0.01 for both). The Ph-based model outperformed the Env-based model using the delta EEG band (WSRT, p = 0.04) and both the Env- and Sgram-models using the theta EEG band (WSRT, p < 0.01). The spectrogram-based model outperformed the envelope-based model using the theta EEG band (WSRT, p = 0.01) but not the delta EEG band. Using the alpha band, the FS model outperformed both the Env model (WSRT, p = 0.01) and the Sgram model (WSRT, p < 0.01). No difference could be observed between the different speech representations using the beta or the low-gamma EEG frequency bands, with correlations below the level of significance (at p = 0.01, significance is at around 0.007).

**Fig. 3.**
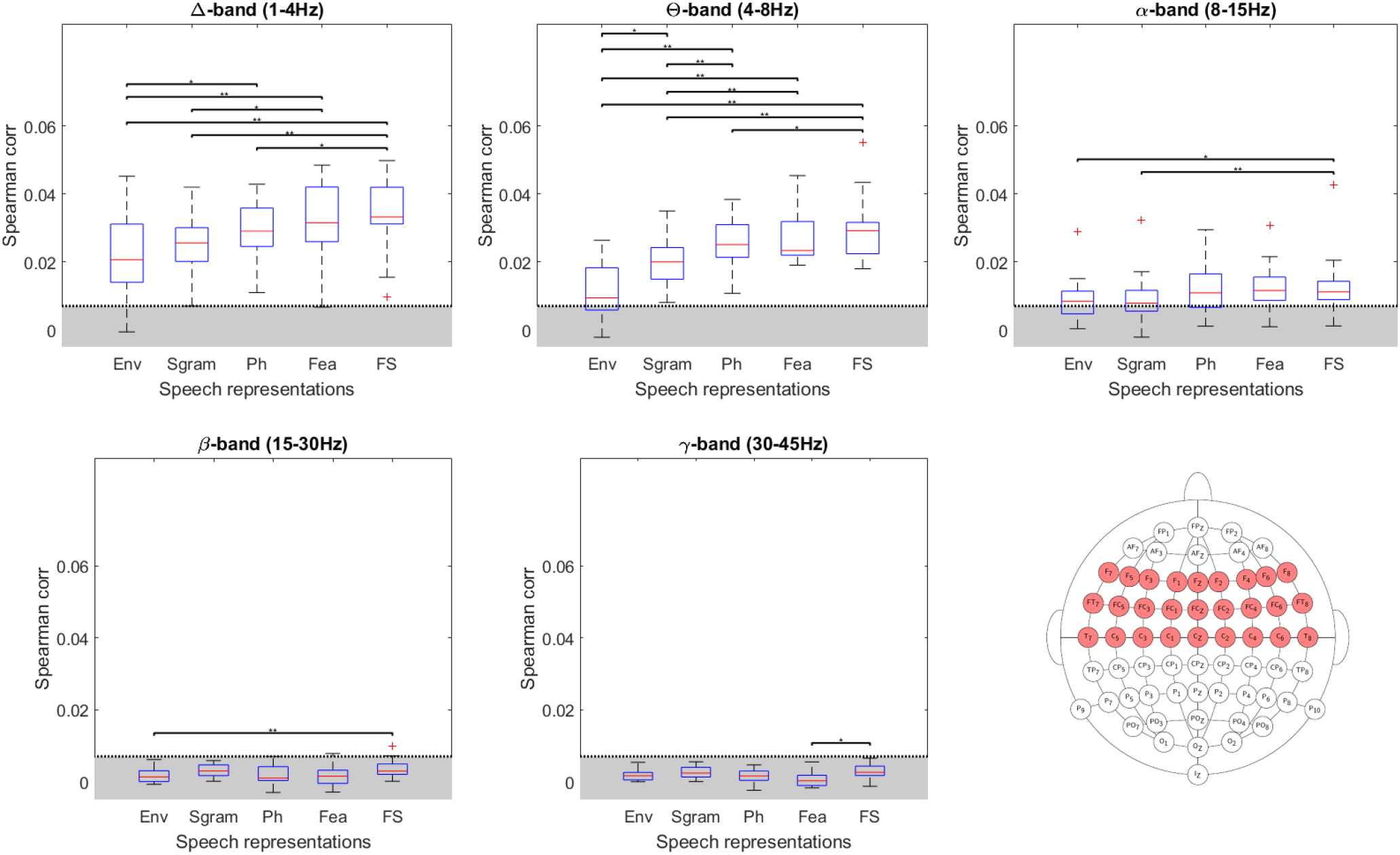
Grand-average EEG prediction correlation (Spearman’s ρ) for each speech model using delta, theta, alpha, beta, and low-gamma EEG frequencies (signal processing scheme B). In grey, the level of significance at p = 0.01. Note the improved EEG prediction using lower frequency bands. As a reminder, for each participant, we computed a subject-specific covariance matrix using six out of the seven folds (corresponding to 12 out of the 14 minutes) of the whole recording (see cross-fold validation). We then computed a grand-average TRF by averaging the covariance matrix across subjects. Finally, this generic model (i.e., the grand-average TRF) was used to predict EEG signals using each stimulus representation from the test set (= the remaining fold, corresponding to two minutes of data). Each participant’s averaged cortical speech tracking was computed as the Spearman’s correlation between the recorded and predicted EEG signals in a selection of fronto-temporal electrodes (see low right panel). The errors bars represents correlation values from the different participants (i.e., 18 data points).

**Fig. 4.**
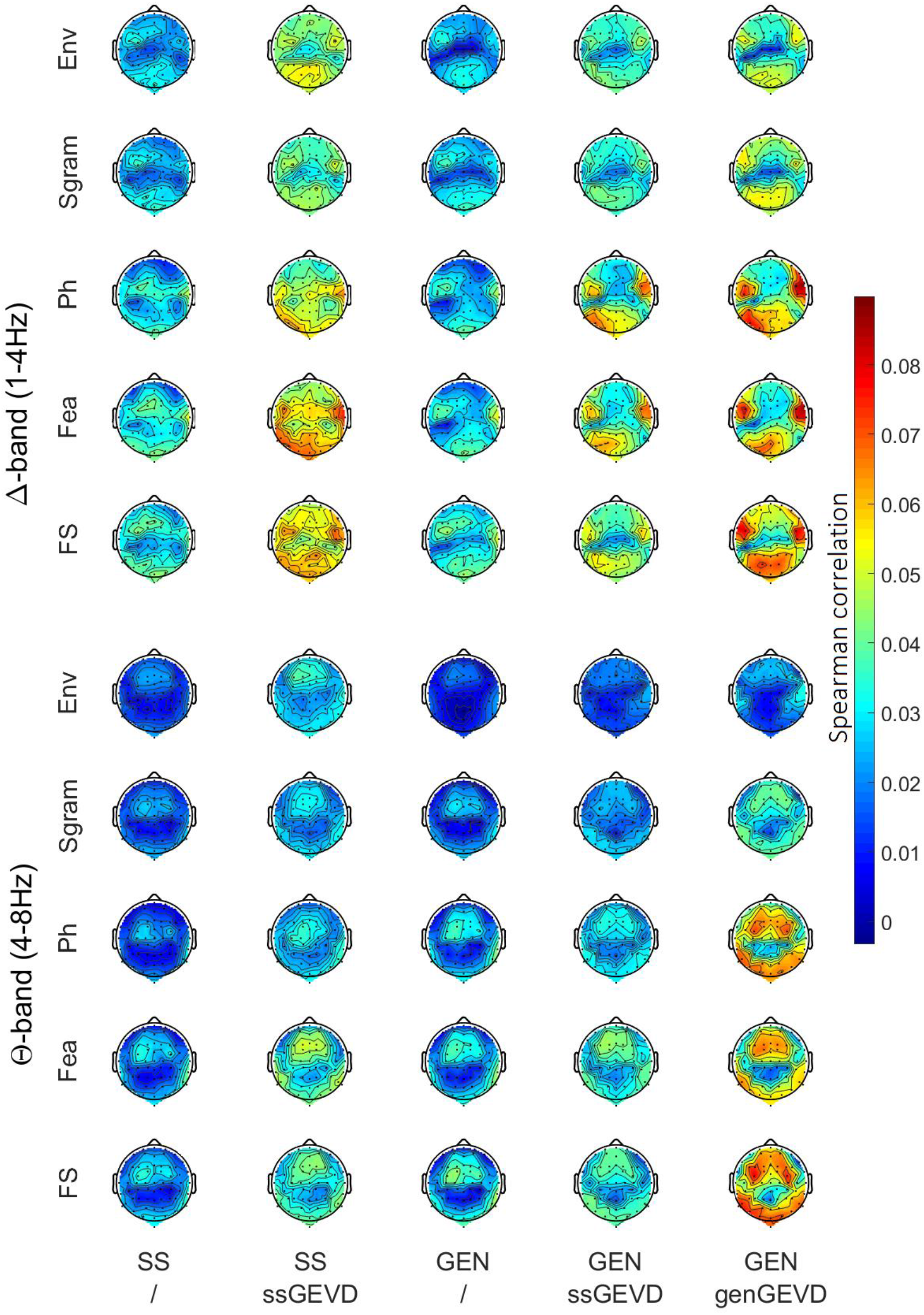
Spatial distribution of the cortical tracking of each speech representation over the scalp (64 electrodes) using subject-specific (ss) or group (gen) data, as well as a GEVD approach (using the five first components). Note the temporal-occipital dominance of the cortical tracking using delta EEG frequency band, while the theta dominance seems more centralized in the frontal-temporal areas.

A visualization of the EEG prediction over the scalp using the different approaches is shown in Fig. 4. For this figure, we used the five first components for the GEVD-based methods. Later in this section, we will show that five components do not yield the highest EEG prediction values (which are obtained using only a few components) but 5 components correspond to a plateau in the EEG predictions without a loss in the spatial resolution (see Fig. 5). Note the increase of the EEG prediction accuracy with the use of GEVD-based methods, and the highlight of the areas of interest (i.e., the spots in the fronto-temporal areas as well as parieto-occipital areas). Interestingly, for the delta EEG band, we observed a temporal and occipital dominance while for the theta EEG band, we observed higher EEG prediction in the centralized frontal area. Focusing on the different ss and gen models, we observed a left delta dominance in the temporo-frontal area for the Env and Sgram model, while the other models presented a wider area of higher correlation connecting the left and right temporal areas. The ss-theta and gen-theta models showed a shared area of activation between the different speech representations; the correlation value of this area increased from Env to FS representations. Looking at the GEVD-based delta models, the left/right balance seems less pronounced and both sides show higher correlations than ss and gen models. The theta GEVD models also showed a centralized dominance better highlighted by the genGEVDgen models.

**Fig. 5.**
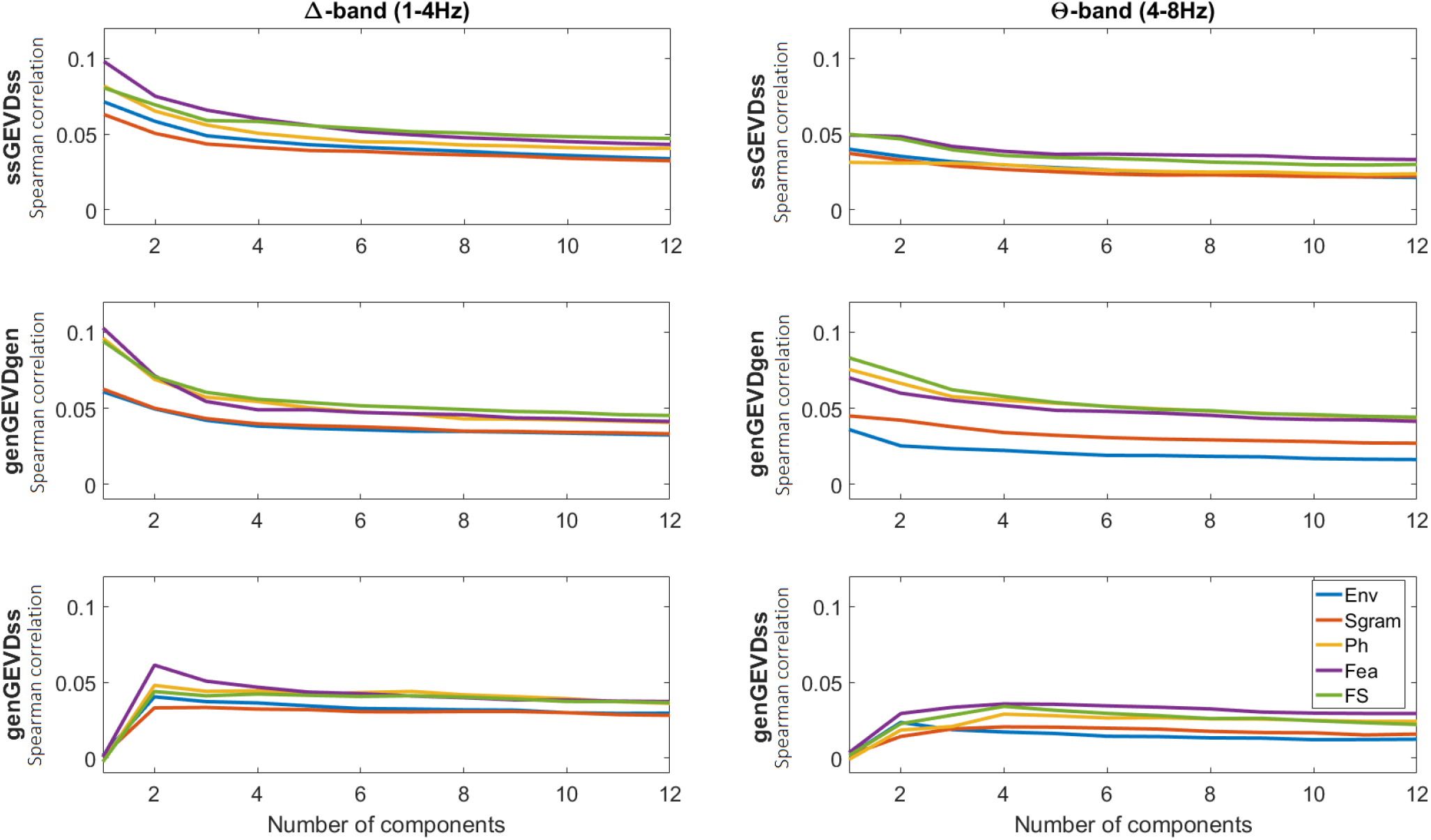
Effect of number of components included in the GEVD-based modeling of the brain responses to the speech. Note the decrease of the averaged Spearman correlation with increasing number of components, except for the hybrid genGEVDss approach.

For both the genGEVDgen and the ssGEVDss models, the highest EEG prediction could be reached using only one GEVD component for either a model based on the delta or theta EEG activity (see Fig. 5, first and second rows). This suggests that stimulus-related activity could be captured by picking the EEG responses associated with the first GEVD component. Predictability decreases with the number of components until reaching a plateau at around five components. The hybrid genGEVDss model showed a maximum correlation using two (resp. four) components using EEG delta (resp. theta) activity (see Fig. 5, third row).

We investigated the addition of a GEVD approach in the different models using one (ssGEVDss and genGEVDgen), two (genGEVDss with the delta band) or four (genGEVDss with the delta band) components (see Fig. 6). In the following, we will mention the difference in the EEG prediction (i.e., the difference in the averaged Spearman correlations over the brain area shown in Fig. 3) between either the ss model and the ssGEVDss model, the ss model and the hybrid genGEVDss model, and finally the gen model and the genGEVDgen model.

**Fig. 6.**
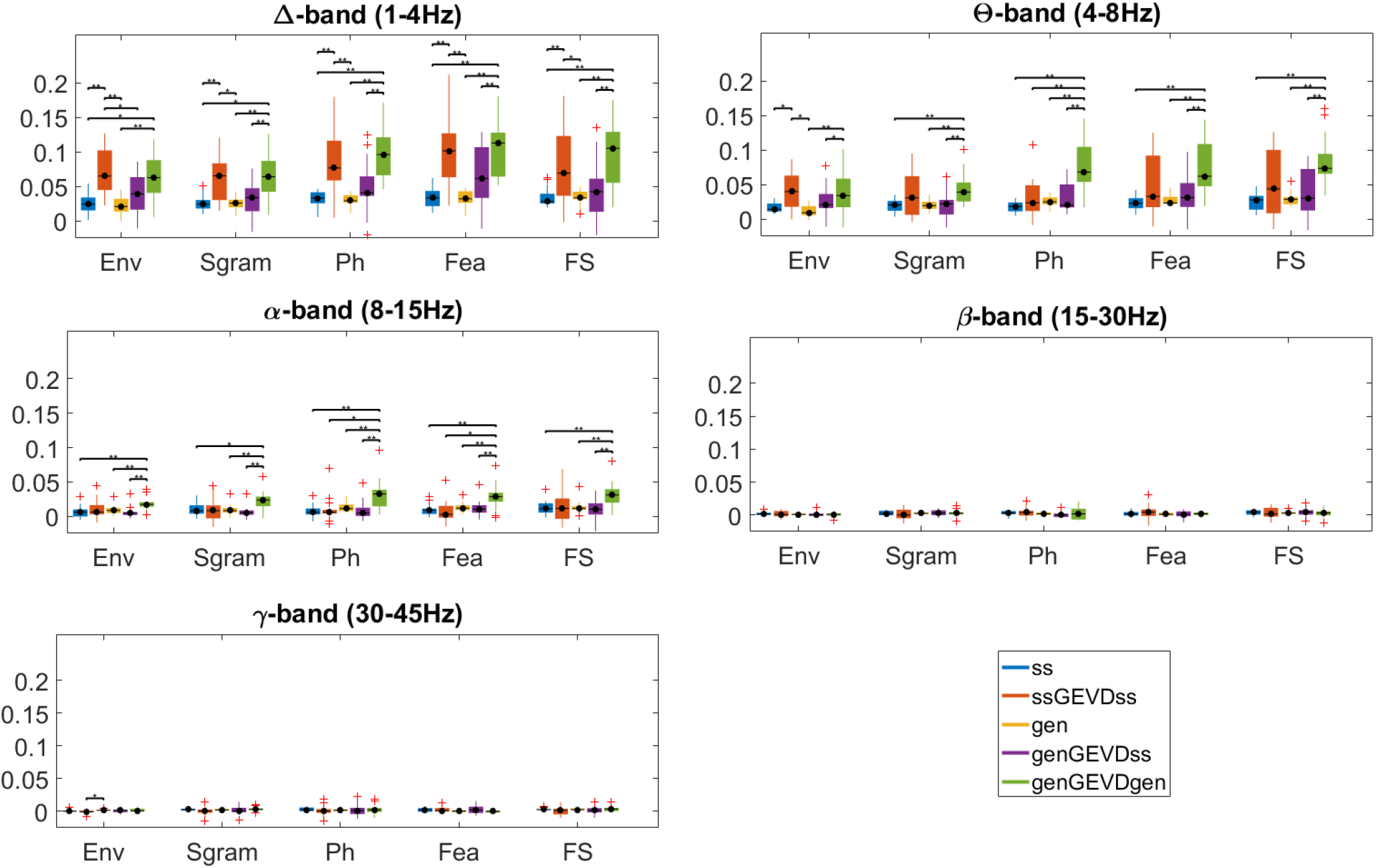
Impact of the inclusion of a GEVD mapping approach on the averaged performance using different EEG frequency bands (delta, theta, alpha, beta, and low gamma) and speech representations (Env, Sgram, Ph, Fea, and FS) as an input to the linear model. Note the improved model using a GEVD approach for both single-subject and group-level decoder. The errors bars represents correlation values from the participants (i.e., 18 data points).

Using the EEG delta band, the addition of GEVD results in an increase in correlation (WSRT, p < 0.01 for all comparisons) for the individual dataset (i.e., ss vs. ssGEVDss) of 0.045 (Env/FS models), 0.050 (Ph), 0.040 (Sgram) and 0.065 (Fea). The maximum Spearman correlation (0.100 ± 0.050) is reached using the ssGEVDss-Fea model. Using the EEG theta band, the addition of a GEVD approach allows increasing the correlation for the Env model by 0.025 (WSRT, p = 0.03). No differences could be observed using higher EEG frequencies in single-subject models (increases in correlation were below 0.005; WSRT, p > 0.05). Interestingly, the hybrid genGEVDss model improved the prediction of EEG responses to none of the speech features (WSRT, p > 0.05).

Using a genGEVDgen model provides significant increases for each model using either the delta, theta or alpha EEG bands (delta: an increase of 0.040 for Env/Sgram, 0.060 for FS, 0.065 for Ph and 0.070 for Fea; theta: an increase of 0.025 for Env/Sgram, 0.055 for FS, 0.050 for Ph and 0.040 for Fea; alpha: an increase of 0.010 for Env/Sgram, 0.020 for FS/Ph and 0.015 for Fea). No differences were observed using beta and low-gamma frequency bands (WSRT, p > 0.05).

Different stimulus representations may be coded in different regions of the brain, and therefore require different spatial filters for optimal reconstruction. We analyzed the influence of the speech feature on the spatial filters obtained from the GEVD, to investigate whether different features require different filters. For the delta EEG frequency band and a model based on a grand-average decoder (see Fig.7, first row, first column), we observed that the cortical tracking with a model based on the Env, the Sgram or the FS speech feature is increased with the use of GEVD-based spatial filtering irrespective of the feature used to build the spatial filter (WSRT, p > 0.05 for each of the Bonferroni-corrected comparisons of the correlation with the maximum correlation reached with the model, see rows in Fig. 7). Inversely, generic models based on Ph or Fea features reached highest cortical tracking using spatial filters built on Ph or Fea features only (WSRT, p < 0.01). For the theta EEG band, higher EEG predictions are reached using spatial filters based on Ph, Fea or FS (see Fig.7, first row, second column). For models based on subject-specific data (see Fig.7, second row), increased EEG predictions are reached using a spatial filter based on FS speech feature (i.e., each model reached a maximum correlation using a FS-based spatial filter; the delta Fea model reached a maximum correlation using a Fea-based spatial filter but no significant difference could be observed using a FS-based spatial filter, WSRT, p = 0.07 uncorrected for multiple comparisons).

**Fig. 7.**
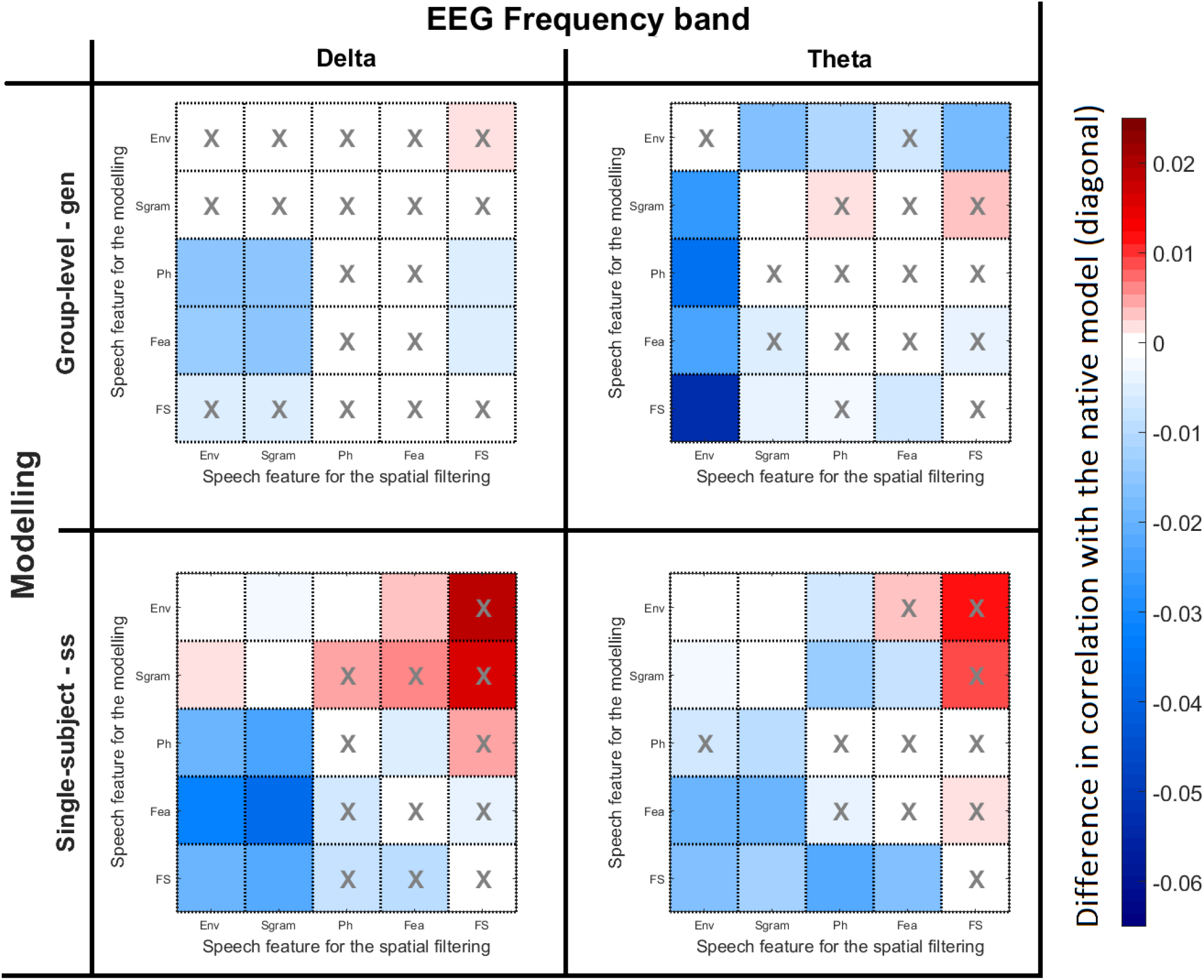
Influence of the speech feature on the spatial filtering of the EEG signals using one GEVD-component, computed as the difference in correlation with the native model. For each model (row), the difference in correlation corresponds to the correlation reached with the spatial filter using the specific speech feature (column) minus the correlation reached using the same speech feature for the model and the spatial filter (see the diagonal). Therefore positive values indicate that the spatial filter under investigation was better than the spatial filter corresponding to the model and vice versa. The y-axis indicates the speech feature used for the modelling of either a generic (first row) or subject-specific (second row) model. The x-axis indicates the speech feature used for computing the spatial filter of the EEG signals over the delta (first column) or theta (second column) EEG signals. For each model (row), a cross means that this spatial filter provided an optimal correlation (e.g., using a generic model and the delta EEG band, a Fea-based model reached an optimal correlation using either a Ph-based spatial filter or a Fea-based spatial filter). Interestingly, spatial filters built on another set of features could also improve the correlation.

## 4. Discussion

The present study illustrates the potential of improving the modeling (TRF) of the brain responses to a speech stimulus by optimally combining EEG channels using a GEVD approach for predicting the brain responses to different speech representations. We showed that the addition of this linear spatial filter allows improving both EEG prediction using subject-specific and generic approaches. In addition, using GEVD avoids manual selection of channels, a limitation of standard forward models.

Consistent with Di Liberto et al (2015), we showed an improvement of the grand-average EEG prediction (gen) using a FS model as compared to the Env, Sgram and Ph speech representations. Each hierarchical stage of the brain encodes a different structure of the speech: the primary stages encodes the acoustical information (e.g., the envelope/spectrogram of the speech) while the later areas encode the discrete tokens of the speech (e.g., phoneme, word, etc). As discussed in Di Liberto et al (2015), the improved EEG predictions highlight the benefit of using both low- and higher-level speech representations to model the brain responses-speech interaction. Interestingly, we also showed that higher representations capture increased information, as suggested by the increases of the cortical tracking to either Ph and Fea, as compared to Env and Sgram. However, we did not find any difference between Fea and FS-based models. Also, while Di Liberto et al (2015) showed an increase of the Pearson’s correlation from the delta to the theta EEG band (Di Liberto et al., 2015), we here observed a decrease of the EEG prediction with increasing EEG frequencies. This could be explained by a difference in the size of the dataset used for training and testing the decoder, or a difference in prosodic features between English and Dutch languages (Moskvina, 2013). Indeed, while Di Liberto recorded EEG in response to an English speech during a story lasting more than one hour, our Dutch-narrated story lasted only around 14 min. They later showed that an averaged difference between the Pearson’s correlations for the FS and S models (in their paper, referred to as FS-S neural index) of around 0.020 could be observed using only 10-min and a grand-average model (Di Liberto & Lalor, 2017) while we here observed a difference of 0.010 between the two models using 12-min of data and a grand-average model. Interestingly, this FS-S neural index increases to 0.04 using the genGEVDgen approach. Note that our electrode selection differs from Di Liberto et al (2015), who only used the six electrodes with the highest averaged correlation over the group in each temporal area. Based on the topographical differences observed between the different frequency bands, we here decided to include central, frontal, as well as temporal electrodes.

Interestingly, the temporal responses of high-level speech showed delayed activations (P1 at around 75 ms, N1 at around 125 ms and P2 at 175ms) as compared to low-level speech representations (P1 at around 50 ms, N1 at around 90 ms and P2 at 140ms). This is in agreement with literature suggesting higher-order brain areas processing phonemes, compared to the acoustic envelope. Note that the long vowels showed a decreased P1 and absent N1 with a strong P2, while other phonetic features showed both a P1 and N1 activations. This could be explained by the difficulty in precisely detecting the onset to a long vowel, resulting in smeared TRFs as compared to other phonetic groups.

The comparison of the topographic distribution of the cortical tracking of the speech over the scalp showed a temporo-occipital dominance in the delta EEG band. We also observed a central-frontal dominance of the EEG prediction to each speech representation in the theta band. Ding & Simon (2014) (Nai Ding & Simon, 2014) suggested that the theta band encodes syllabic information (Drullman, Festen, & Plomp, 1994b, 1994a; Elliott & Theunissen, 2009) present in the envelope while the delta EEG band is sensitive to the envelope prosody, in particular the sentence and word rhythms (Goswami & Leong, 2013). We here hypothesized that the frontal area, associated with top-down control, is more involved in the encoding of syllabic features critical for speech recognition, explaining this frontal-theta dominance, while early brain areas processed the encoding of prosodic information of speech, linked with the delta cortical tracking of the speech.

According to our results, the GEVD-based spatial filter improves higher-level models more than low-level models. We can make several hypotheses: (1) the spatial filter and the brain-stimulus model positively reinforce each other, i.e., a good model allows to optimally predict the EEG responses to speech, allowing a good spatial filter estimate, resulting in an even better mapping between the actual and the predicted EEG after spatial filtering; (2) higher-level speech features activate a broader network than the low-level ones, thus a robust mapping can more easily be extracted from each individual.

The application of a GEVD-based spatial filter on either subject-specific or group data improved the EEG prediction accuracy for all the different speech features, using only one component. No difference could be observed between a fully generic or full subject-specific approach (i.e., genGEVDgen and ssGEVDss). However, the hybrid genGEVDss showed significantly lower cortical tracking using one brain component (see Fig. 6). It is important to note that a generic model will focus on cortical responses that are consistent across subjects while a subject-specific model will take into account anatomical and physiological differences between individuals. Consequently, we can here hypothesize that a genGEVDss approach will first spatially filter the EEG responses to find components that generalize between subjects and will then look for specific responses, which are probably removed from the EEG signal if only one generalized component is used.

To investigate whether different speech features require specific spatial filters, we evaluated the impact of a spatial filter built on a specific speech feature on a model based on a different speech feature. In the case of a generic model, we observed that model based on low-level speech features could benefit from a spatial filter built on either (or both) low-level and higher-level speech features when using the delta EEG frequencies. Inversely, models based on higher-level speech features required higher-level speech features to reach optimal EEG predictions. We hypothesize that higher-level speech features are coded by a broader brain network, that also incorporates regions that code lower-level features. For models based on single-subject data, spatial filters based on FS provides the best EEG predictions for all models. We hypothesize that subject-specific models benefit from the positive reinforcement effect due to generally improved performance of the FS model mentioned above.

Our pipeline consists of two different data processing stages: the modelling of the brain-speech interaction and the spatial filtering. The model requires more training data than the spatial filter, so the spatial filter does not require additional data. Consequently, we required around 10-min of data in order to properly model the brain-speech interaction and therefore ensure a proper selection of the spatial filter’s components (reminder, we first need to predict the EEG from the training dataset, then map it to the recorded EEG). A substantial decrease in the performance has been shown in both Auditory-Attention and Cortical-Tracking protocols when dealing with models based on less than 10-min of training data (Di Liberto & Lalor, 2017; Mirkovic, Debener, Jaeger, & De Vos, 2015).

We demonstrated that the brain responses to naturalistic speech could be better predicted by including a phoneme-based stimulus representation, and by extracting speech-related information from the recorded EEG signals using a GEVD approach combined with standard multivariate TRF (Crosse et al., 2016). This method, which is closely related to the CCA-based method proposed by de Cheveigné & Parra (2014) and Dmochowski, Ki, DeGuzman, Sajda, & Parra (2018), allows substantially increasing the EEG prediction, with a correlation of around 0.1 reached using the delta EEG band and a high-level speech representation. Moreover, we here showed that the inclusion of this linear spatial mapping allows removing the impact of manual channel selection on the performance by automatically weighting channels in a data-driven way. Future research should evaluate the inclusion of higher-level speech features such as words or semantic information, phoneme-level surprisal, entropy, or phoneme onset in the FS-model (Brodbeck, Hong, & Simon, 2018; Broderick, Anderson, Di Liberto, Crosse, & Lalor, 2018).

## Acknowledgements

The authors would like to thank Giovanni Di Liberto for his helpful advice on the analysis, as well as Hugo Van hamme and Lien Decruy for helping with the phoneme segmentation. Financial support was provided by the KU Leuven Special Research Fund under grant OT/14/119 to Tom Francart. This project has received funding from the European Research Council (ERC) under the European Union’s Horizon 2020 research and innovation program (grant agreement No 637424, ERC starting Grant to Tom Francart). Research funded by a PhD grant of the Research Foundation Flanders (FWO) for Jonas Vanthornhout (1S10416N) and Eline Verschueren (1S86118N). The authors declare that they have no conflict of interest

